# LRRC23 is a conserved component of the radial spoke that is necessary for sperm motility and male fertility in mice

**DOI:** 10.1101/2021.06.01.446532

**Authors:** Xin Zhang, Jiang Sun, Yonggang Lu, Jintao Zhang, Keisuke Shimada, Taichi Noda, Shuqin Zhao, Takayuki Koyano, Makoto Matsuyama, Shushu Zhou, Jiayan Wu, Masahito Ikawa, Mingxi Liu

## Abstract

Cilia and flagella are ancient structures that achieve controlled motor functions through the coordinated interaction of structures including dynein arms, radial spokes (RSs), microtubules, and the dynein regulatory complex (DRC). RSs facilitate the beating motion of these organelles by mediating signal transduction between dyneins and a central pair (CP) of singlet microtubules. RS complex isolation from *Chlamydomonas* axonemes enabled the detection of 23 different proteins (RSP1-23), with the roles of RSP13, RSP15, RSP18, RSP19, and RSP21 remained poorly understood. Herein, we show that *Lrrc23* is an evolutionarily conserved testis-enriched gene encoding an RSP15 homolog in mice. Through immunoelectron microscopy, we demonstrate that LRRC23 localizes to the RS complex within murine sperm flagella. We further found that LRRC23 was able to interact with RSHP9 and RSPH3A/B. The knockout of *Lrrc23* resulted in RS disorganization and impaired motility in murine spermatozoa, whereas the ciliary beating was unaffected by the loss of this protein. Spermatozoa lacking LRRC23 were unable to efficiently pass through the uterotubal junction and exhibited defective zona penetration. Together these data indicate that LRRC23 is a key regulator underpinning the integrity of RS complex within the flagella of mammalian spermatozoa, whereas it is dispensable in cilia.

**Author summary:** 

## Introduction

Flagella are essential mediators of male gamete motility in eukaryotic species. In mammals, the sperm flagella are characterized by the middle, principle, and end segments through which the axonemal structure runs [1]. Most prior studies of axonemal motility focused on sea urchin spermatozoa or on ciliated/flagellar unicellular organisms to gain insight into mammalian sperm functionality. Dynein motors serve as essential mediators of axonemal motility, and are composed of outer and inner arms lining along the longitudinal axis of the axonemal cylinder. These axonemal dyneins exhibit ATP-insensitive anchoring to the A-tubule in each doublet, whereas they undergo ATP-dependent stepping motion along the neighboring doublet B-tubules, resulting in sliding between these doublets. Restriction of such sliding by structures including nexin links, converts such motion into axonemal bending [2, 3]. A central pair (CP) of singlet microtubules, radial spokes (RSs), the I1 inner arm dynein (IDA), and the dynein regulatory complex (DRC) are essential mediators of dynein motility within this context [4–8]. Signal from the CP complex is passed through RSs to the IDAs, and the I1 dynein intermediate chain–light chain (IC–LC) complex and the DRC signal through IDAs and outer dynein–inner dynein (OID) linkers to outer dynein arms (ODAs) [9]. In addition to mediate the beating motion of flagella by relaying signals between the CP and dynein proteins [10], the RS complex is also important for maintaining the stability of the ‘9 + 2’ axonemal structure [11, 12]. Through studies on *Chlamydomonas reinhardtii*, purification of the RS complex led to the identification of 23 *Chlamydomonas* flagellar RS proteins (RSP1-23) [13–16].

While RS complexes exhibit a similar T-shaped morphological orientation across species, RS evolutionary divergence has been noted, with many evolutionary changes having favored the simplification of this multi-protein complex [17]. In line with the predicted partial redundancy of spoke head proteins, RSP1 and its putative binding partner NDK5, for example, are not present in *Tetrahymena* or other ciliates, while RSP4/6 are encoded by a single gene in *Ciona intestinalis* and sea urchins [17]. Indeed, only one RSP4/6 protein and one MORN protein are detected in purified *Ciona* RS samples [18]. Similar changes may have also influenced RS complex development in humans, although humans do possess orthologs of *RSP1*, *4*, *6*, and *10*. The expression of some of these genes is tissue-specific in humans, with *RSP1* (*RSPH1*) and *RSPH10* exhibiting inverse expression patterns in airway and testis tissues [19]. We have recently demonstrated that RSPH6A is enriched in murine testis wherein it localizes to the sperm flagella. Male *Rsph6a* knockout mice exhibit infertility attributable to their short immotile sperm flagella [11]. Overall, these data suggest that RS structures in mammals are distinct from those in other species and differ between cilia and flagella. However, the latter differences emphasize the importance of conducting further experiments to verify the localizations and functions of different RS component proteins in mammalian species.

In *Chlamydomonas*, these RSPs are assembled to yield the RS complex in two primary stages. Cell body extract fractionation experiments have revealed that a 12S RS precursor complex is assembled in the cell body [20]. Intraflagellar transport facilitates the delivery of these 12S precursors into flagella wherein they undergo conversion to yield the mature 20S RS complex. RSP1-7 and 9-12 compose the 12S precursor complex, with the remaining RSPs being assembled following transportation to the axoneme [21]. Nevertheless, the specific roles of RSP13, 15, 18, 19, and 21 in the maturely assembled RS complex are less understood. *Chlamydomonas* RSP15 is known to be a leucine-rich repeat (LRR) protein that is thought to be homologous to LRR37 protein found within the RSs in the spermatozoa of *Ciona intestinalis* [14, 22]. Mutations in the LRR37 homolog LRRC23 result in defective ciliary motility in the cilia of the otic vesicle in zebrafish [23]. Mutations in LRRC23 also cause defective phagocytosis and reduced swimming velocity in *Tetrahymena*, consistent with the ciliary defects [23]. The functional role of LRRC23 in mammals, however, remains to be evaluated.

Herein, we show that *Lrrc23* is an evolutionarily conserved gene preferentially enriched in testis and required for male fertility in mice. LRRC23 localized to the RS and interacted with RSHP9 and RSPH3A/B, and the knockout of *Lrrc23* resulted in RS complex disorganization and impaired murine sperm motility, whereas RS distribution and motility in cilia from these mice was unaffected. Together, these data suggest that LRRC23 plays an essential role in stabilizing the RS complex in sperm flagella, but is dispensable in respiratory cilia.

## Results

### *Lrrc23* is an evolutionarily conserved gene enriched in the testis

*Lrrc23* is encoded by the genomes of known basal eukaryotic species that utilize flagella during at least one life cycle stage (S1A Fig). In humans, the LRRC23 protein contains eight leucine-rich repeat (LRR) domains and a coiled-coil domain in its N-terminal region, with the LRRs of this protein being conserved among species (S1B Fig). We began the present study by profiling *Lrrc23* tissue-specific expression patterns via RT-PCR in adult mice, revealing a distinct band in the testis and a weaker band in lung tissues (Fig 1A). We then evaluated the expression of *Lrrc23* in postnatal testis to follow the leading edge of the first wave of spermatogenesis. This analysis revealed *Lrrc23* expression initiated on postnatal day 14, which is roughly consistent with the first appearance of pachytene spermatocytes (Fig 1B). Similar trends were observed when reviewing published single-cell RNA-seq data pertaining to spermatogenesis in humans and mice (S2 Fig) [24].

**Fig 1.**
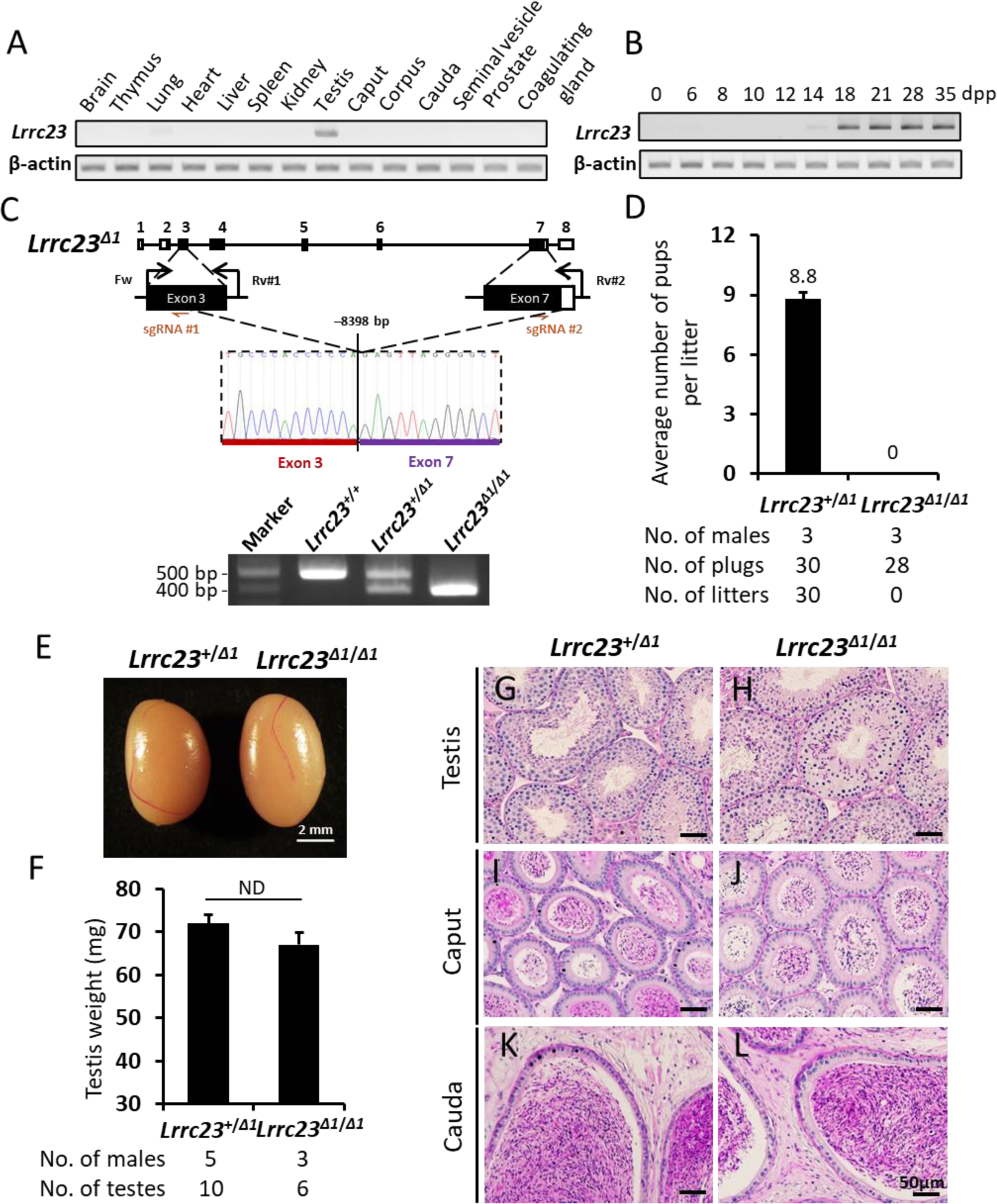
Generation and analysis of male *Lrrc23^Δ1/Δ1^* mice. (A) Murine *Lrrc23* expression in the indicated organs was assessed via RT-PCR, with *Actb* serving as a control. (B) Murine *Lrrc23* expression in the testis tissue samples collected on the indicated day postpartum (dpp) was analyzed via RT-PCR with *Actb* as a normalization control.(C) The genomic structure of *Lrrc23* and the CRISPR/Cas9 targeting approach. Dual sgRNAs (sgRNA#1 and sgRNA#2) were respectively used to target exons 3 and 7. The deletion of an 8398 bp fragment of *Lrrc23* between exons 3 and 7 was confirmed via Sanger sequencing and PCR. The coding region is indicated using black rectangles. Genotyping primers (Fw, Rv#1, and Rv#2) were as shown. (D) Numbers of pups born per vaginal plug detected in the indicated groups. N = 3 males each for *Lrrc23^+/Δ1^* and *Lrrc23^Δ1/Δ1^* mice, *P* < 0.05. (E) Testes of *Lrrc23^+/Δ1^* and *Lrrc23^Δ1/Δ1^* mice. (F) A comparison of the weights of testes from *Lrrc23^+/Δ1^* (N = 5) and *Lrrc23^Δ1/Δ1^* (N = 3) mice, *P* > 0.05. (G-L) Testes and epididymides from *Lrrc23^+/Δ1^* and *Lrrc23^Δ1/Δ1^* mice were subjected to histological staining.

### *Lrrc23* is essential for male fertility and sperm motility

To test the functional importance of *Lrrc23* in mice, we employed the CRISPR/Cas9 genome editing system to generate two *Lrrc23* mutant mouse lines. We first prepared a stable *Lrrc23* mutant mouse line harboring a deletion of the exons 3-7 of this gene (*Lrrc23^Δ1/Δ1^*; Fig 1C). Male *Lrrc23^Δ1/Δ1^* mice did not exhibit any overt developmental or behavioral abnormalities. We assessed their fertility by housing individual *Lrrc23^+/Δ1^* and *Lrrc23^Δ1/Δ1^* males with wildtype (WT) females and counting the numbers of offspring. *Lrrc23^Δ1/Δ1^* males were unable to sire any offspring despite successful copulation with WT females confirmed by vaginal plugs (Fig 1D), suggesting the loss of *Lrrc23* expression results in male infertility in mice.

In order to determine whether the male infertility was due to arrested spermatogenesis, we next investigated the sperm formation and production in *Lrrc23^Δ1/Δ1^* male animals. However, no differences in testis appearance and weight were detected when comparing *Lrrc23^+/Δ1^* and *Lrrc23^Δ1/Δ1^* littermates (Fig 1E and 1F). Likewise, periodic acid and Schiff’s reagent (PAS) staining and counterstaining with Mayer hematoxylin solution of testicular sections failed to reveal any differences in the spermatogenesis between *Lrrc23^+/Δ1^* and *Lrrc23^Δ1/Δ1^* males (Fig 1G and 1H). Epididymal ducts filled with spermatozoa were observed in both the cauda and caput regions (Fig 1I-1L).

To further determine the cause of male infertility in *Lrrc23* knockout mice, we conducted an in vivo fertilization assay. No fertilized eggs were found in *Lrrc23^+/+^* females after mating with *Lrrc23^Δ1/Δ1^* males, suggesting the male sterility originated from impaired fertilization instead of defective embryogenesis (Fig 2A and 2B). To investigate if the impaired fertilization was derived from problems in sperm migration or downstream zona pellucida (ZP) penetration or gamete fusion, we then carried out a uterotubal junction (UTJ) penetration assay. *Lrrc23* knockout spermatozoa exhibited inability to pass through the UTJ (Fig 2C). Computer-Assisted Sperm Analysis (CASA) was further carried out to assess the sperm motility in *Lrrc23^Δ1/Δ1^* males, revealing a significant reduction in motile spermatozoa and spermatozoa exhibiting progressive motility, a condition known as asthenozoospermia (Fig 2D and 2E). While progressive motility was impaired, the flagella of spermatozoa from *Lrrc23^Δ1/Δ1^* mice did beat, albeit across a more limited range (Fig 2F, S1 Video, S2 Video). We additionally conducted in vitro fertilization (IVF) and determined that *Lrrc23* knockout spermatozoa were unable to fertilize cumulus-intact or cumulus-free ZP-intact oocytes (S3A and S3B Fig). However, following ZP removal, *Lrrc23* knockout spermatozoa were able to fuse with oocytes (S3C Fig), indicating that *Lrrc23* knockout spermatozoa were defective in passing through the UTJ and penetrating the cumulus cell layer and/or the ZP due to impaired motility.

**Fig 2.**
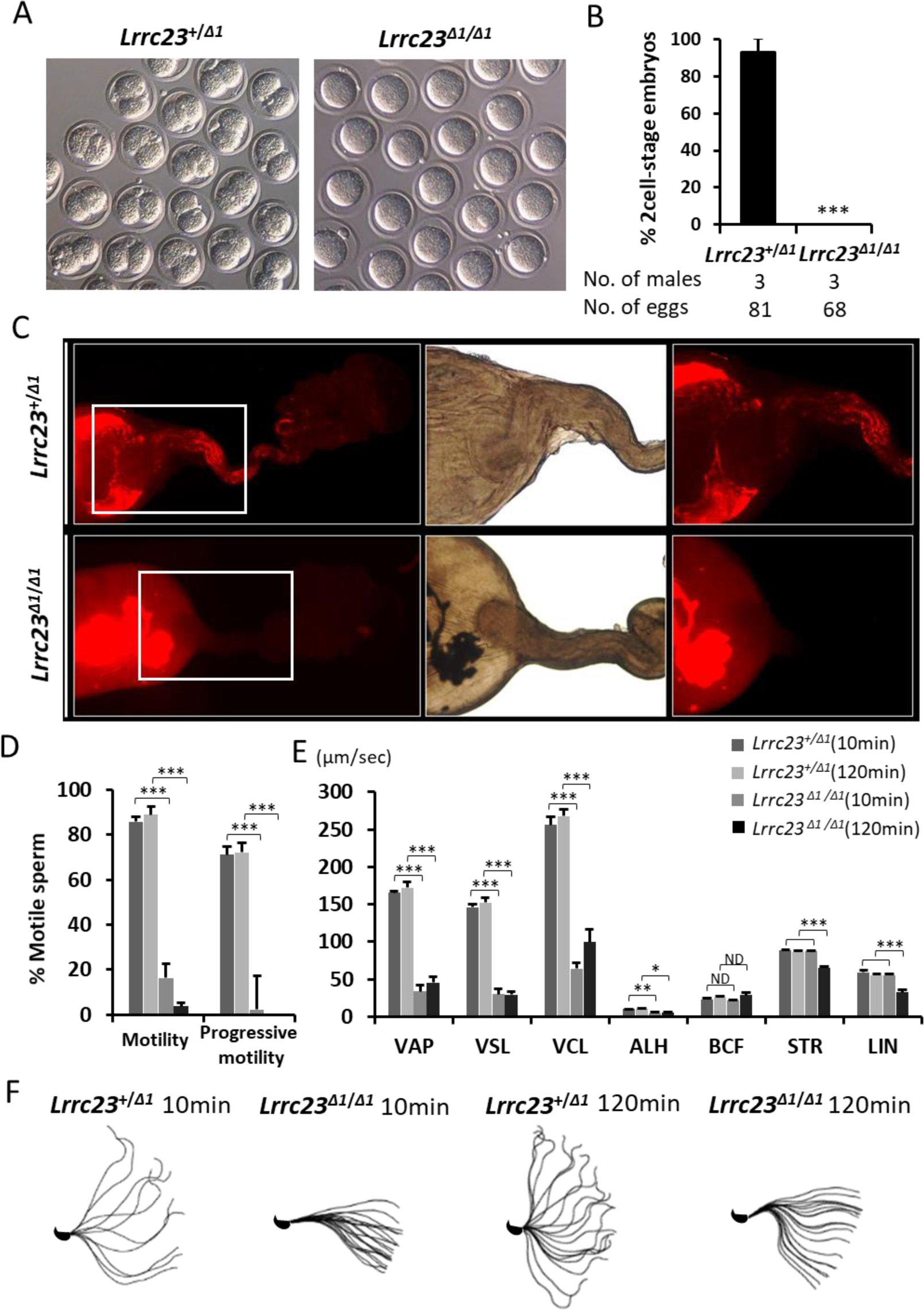
Assessment of the motility and fertility of spermatozoa from Lrrc23^Δ1/Δ1^mice. Images of eggs retrieved from the oviducts of WT females mated with *Lrrc23^+/Δ1^* and *Lrrc23^Δ1/Δ1^* mice during in vivo fertilization test. (B) Percentages of two-cell embryos in cumulus-intact oocytes inseminated with spermatozoa from *Lrrc23^+/Δ1^* and *Lrrc23^Δ1/Δ1^* mice, N = 3, *P* < 0.05. (C) Images of sperm migration through the female reproductive tract. WT females were mated with *Lrrc23^+/Δ1^* and *Lrrc23^Δ1/Δ1^* mice carrying the RBGS transgene such that spermatozoa were fluorescently tagged, revealing the failure of *Lrrc23* knockout spermatozoa to pass through the UTJ. White rectangles indicate magnified regions.(D) Relative percentages of motile and progressively motile sperm from *Lrrc23^+/Δ1^* and *Lrrc23^Δ1/Δ1^* mice after 10 and 120 min of incubation in the TYH medium. (E) Different motility parameters for sperm from *Lrrc23^+/Δ1^* and *Lrrc23^Δ1/Δ1^* mice as determined via CASA following incubation for 10 or 120 min in TYH medium. VCL, curvilinear velocity; VAP, average path velocity; ALH, amplitude of lateral head; VSL, straight-line velocity; STR, straightness; and LIN, linearity. (F) Flagellar waveforms were assessed after incubation for 10 and 120 min, with motility being imaged at 200 frames/second and with individual frames from a single beating cycle being superimposed.

To confirm these observations, we generated another stable *Lrrc23* mutant mouse line harboring a 49 bp deletion in the exon 4 (*Lrrc23^Δ2/Δ2^*; S4A Fig). Likewise, gross examinations revealed no significant differences in testis weight or sperm counts when comparing *Lrrc23^Δ2/Δ2^* and *Lrrc23^+/+^* males (S4B-S4E Fig) and *Lrrc23^Δ2/Δ2^* male mice also exhibited infertility (S4F Fig). No abnormality in sperm morphology was detected in the *Lrrc23^Δ2/Δ2^* males under light microscope (S4G Fig) and scanning electron microscope (SEM; S4H Fig). Consistent with the spermatozoa from *Lrrc23^Δ1/Δ1^* males, CASA revealed a reduction in sperm motility and progressive movement in *Lrrc23^Δ2/Δ2^* mice (S4I-S4K Fig). In summary, these findings together confirm that *Lrrc23* regulates sperm motility and is required for male fertility in mice.

### LRRC23 is a radial spoke component that localizes to sperm flagella

We assessed the subcellular localization of LRRC23 using internally generated antibodies. Western blotting revealed LRRC23 was approximately 40 kDa in size and was absent in *Lrrc23^Δ2/Δ2^* mice (Fig 3A). Immunoblot analyses of sperm protein extracts indicated that LRRC23 was present within the Triton X-100-resistant, SDS-soluble fraction (Fig 3B), which is the fraction associated with the axoneme [25]. We next conducted immunoprecipitation (IP)-mass spectrometry (MS) to characterize proteins interacting with LRRC23 in murine testis, leading to the identification of five unique proteins that were absent only in the control non-immune serum precipitated samples (S5A Fig and Table 1). Among these putative LRRC23-interacting proteins, the RS head component RSPH9 was detected, which was further confirmed by co-immunoprecipitation and Western blot analysis (Fig 3C).

**Fig 3.**
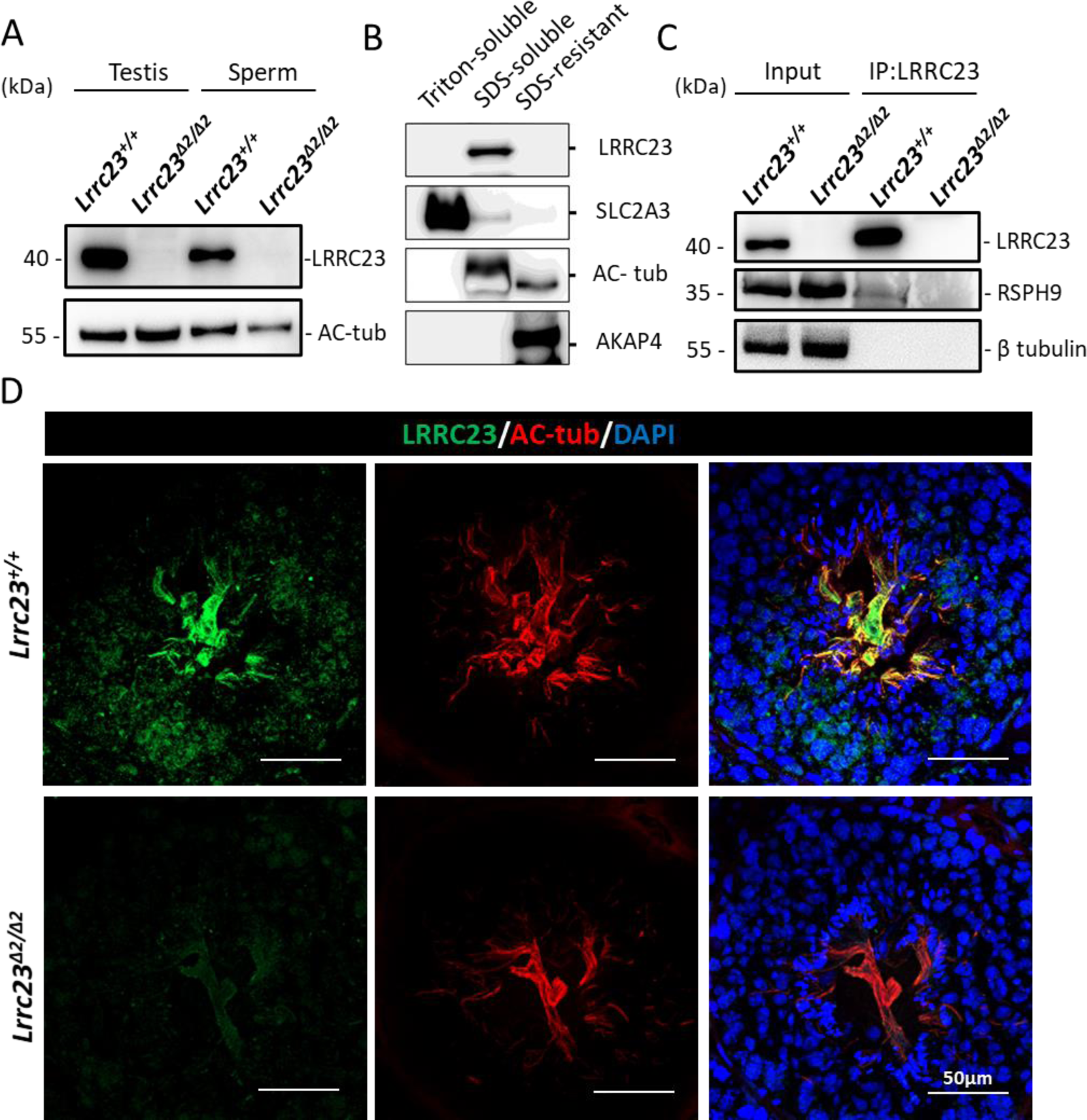
LRRC23 localizes to the sperm tails. (A) LRRC23 expression in the testis and cauda epididymal spermatozoa from *Lrrc23^+/+^* and *Lrrc23^Δ2/Δ2^* mice. No difference in the acetylated tubulin (Ac-tub) staining was detected between the two groups of mice. (B) Murine sperm fractionation revealed the presence of LRRC23 in the SDS-soluble fraction. SLC2A3, acetylated tubulin, and AKAP4 were used to respectively mark the fractions that were Triton-soluble, SDS-soluble, and SDS-resistant. (C) An assessment of LRRC23 and RSPH9 co-IP in testicular protein extracts from *Lrrc23^+/+^* and *Lrrc23^Δ2/Δ2^* mice. (D) Testis cross-sections from WT and *Lrrc23^Δ2/Δ2^* mice were stained with immunofluorescent antibodies specific for AC-tub (red) and LRRC23 (green), with Hoechst 33342 (blue) being used to detect nuclei.

**Table 1.**
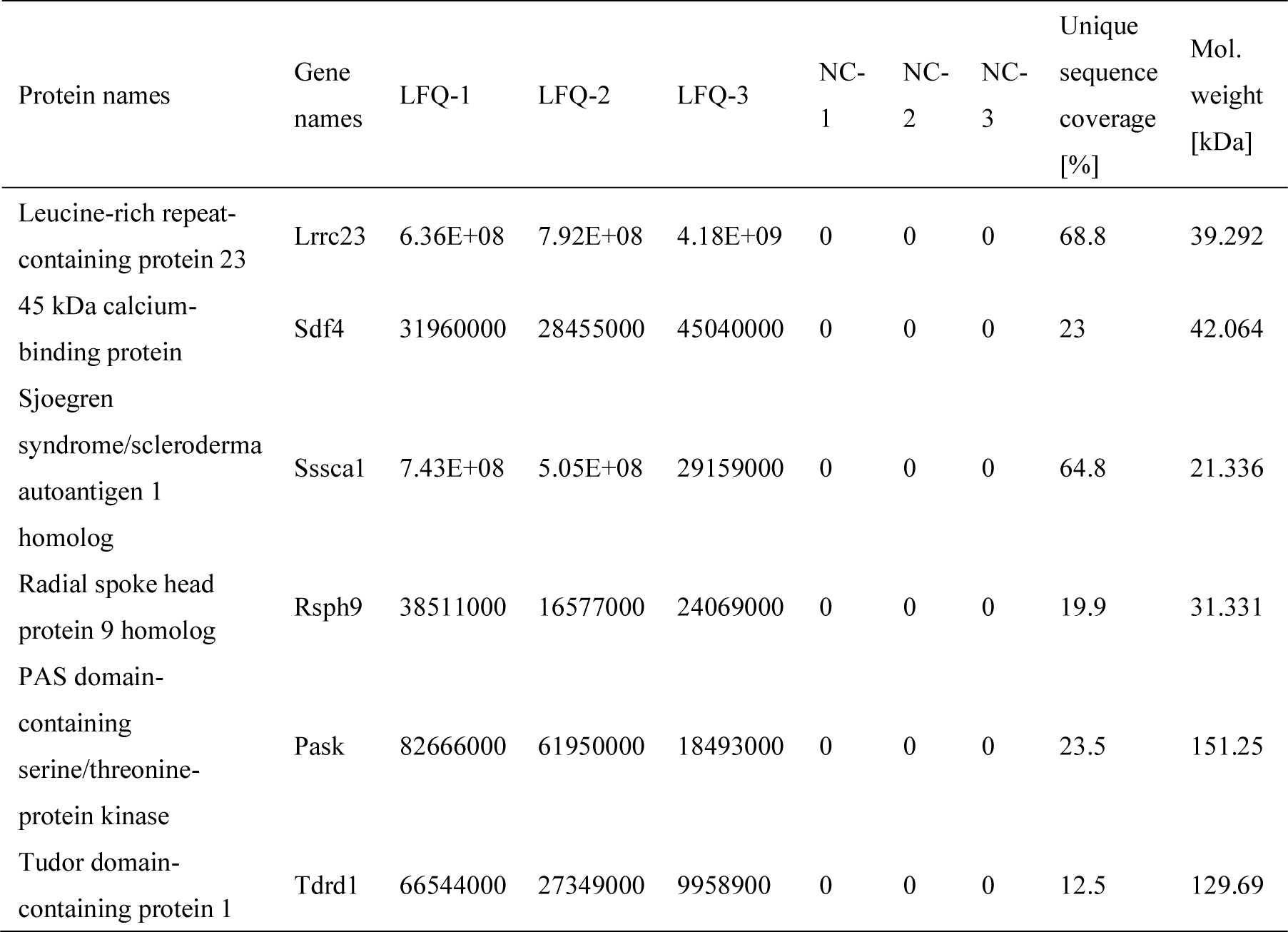
Anlysis of LRRC23 immunoprecipitation (IP) in testicular protein extracts via mass spectrometry (MS).

Confocal microscopic observation revealed LRRC23 presented within the flagella and cytoplasm in murine testicular spermatids (Fig 3D). High-resolution microscopy confirmed LRRC23 localized to the flagella of murine spermatozoa, and was closer to the CP than the acetylated tubulin (Fig 4A). Immunoelectron microscopy uncovered that LRRC23 localized to the RS within spermatozoa (Fig 4B-4K). The LRRC23 homolog RSP15 in *Chlamydomonas* has previously been shown to interact with RSP3 and RSP22 [26]. Although we did not detect any interaction between LRRC23 and RSPH22, we did find that it was able to interact with RSPH3A/B (S5B-S5D Fig), strongly suggesting that LRRC23 is a RS component within sperm flagella.

**Fig 4.**
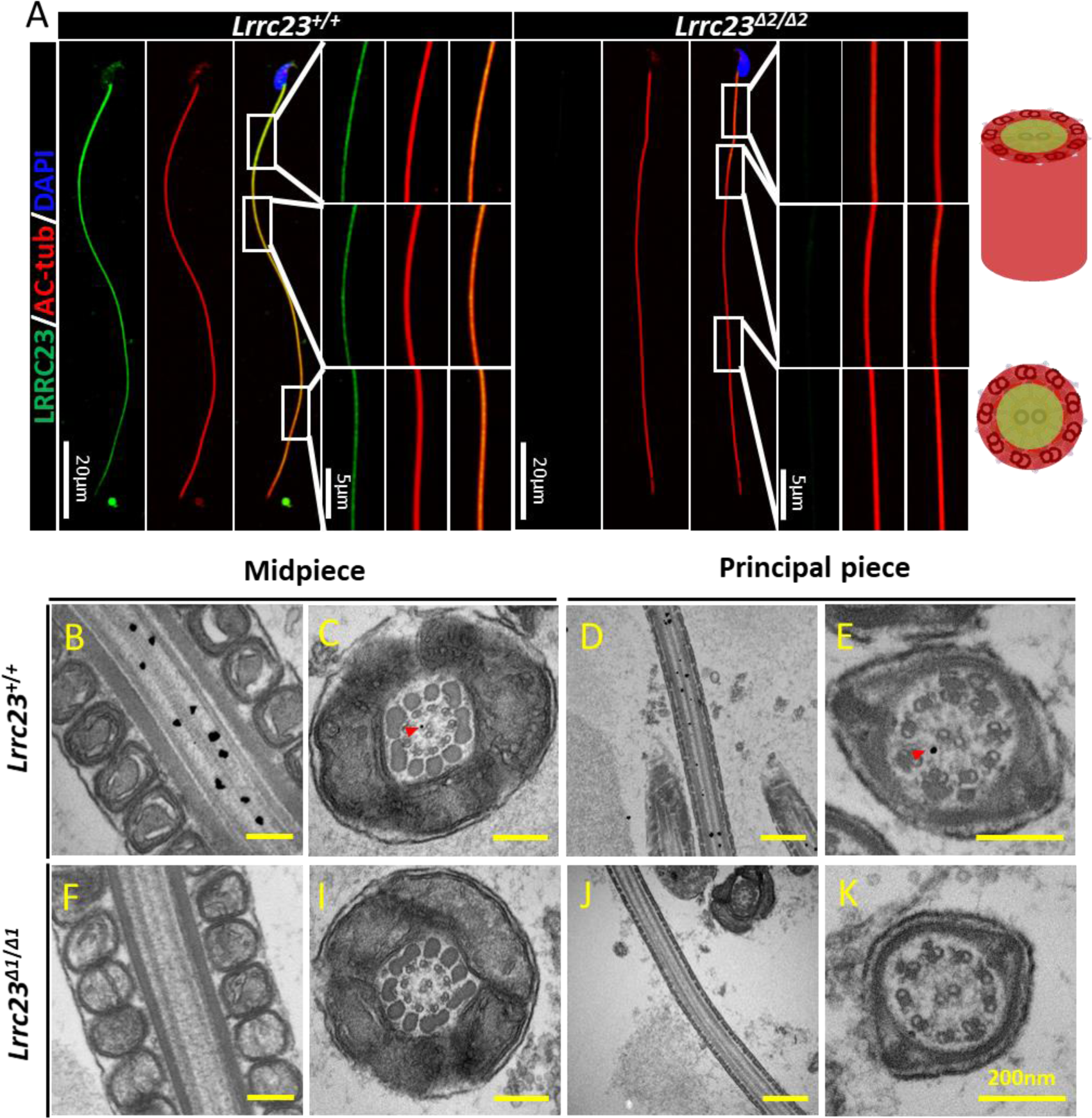
LRRC23 localizes to axonemal radial spokes within sperm flagella. (A) Spermatozoa from *Lrrc23^+/+^* and *Lrrc23^Δ2/Δ2^* mice were subjected to immunofluorescent staining using antibodies specific for AC-tub (red) and LRRC23 (green), with Hoechst 33342 (blue) being used to detect nuclei. AC-tub axonemal localization was evident in the sperm of both WT and *Lrrc23^Δ2/Δ2^* mice. Magnified regions are marked with white rectangles, and panels on the right demonstrate LRRC23 localization to the center of these axonemes. (B-K) Immunoelectron microscopy was conducted to evaluate sperm flagella from *Lrrc23^+/+^* and *Lrrc23^Δ1/Δ1^* mice; (B and D) Longitudinal and (C and E) cross sections of the midpiece and principal piece labeled with anti-LRRC23. Gold particles (red arrowheads) were found to localize within the axoneme between the CP and the outer doublet microtubules consistent with the location of the radial spokes. (F, I, J and K) Sperm flagella from *Lrrc23^Δ1/Δ1^* mice served as controls.

IP-MS also revealed the ability of PAS domain-containing serine/threonine-protein kinase (PASK) to interact with LRRC23, and such an interaction was confirmed in a co-IP experiment (S6A Fig). Western blotting demonstrated that the LRRC23 bands in sperm and testis samples shifted to a lower position after treatment with calf-intestinal alkaline phosphatase (CIP), suggesting that LRRC23 was phosphorylated in vivo (S6B Fig). When cauda epididymal spermatozoa were treated with the protein kinase A (PKA) and PASK inhibitors H-89 and BioE-1115, respectively, we found that neither of these inhibitors significantly affected the LRRC23 band position upon Western blotting (S6C Fig). The treatment of BioE-1115 showed a minor impact on sperm motility at a concentration of 100 μM (S6D and S6E Fig). Together, these data indicated that the post-translational modification of LRRC23 is likely completed during spermatogenesis and that PASK does not regulate the phosphorylation of LRRC23 in mature spermatozoa.

### *Lrrc23* knockout causes abnormal RS formation in sperm flagella but does not adversely impact respiratory cilia

To test the impact of LRRC23 depletion on RS assembly, we next examined the ultrastructure of RSs in the spermatozoa of *Lrrc23^Δ1/Δ1^* males via transmission electron microscopy (TEM), revealing partial RS absence or disorder in the spermatozoa from *Lrrc23^Δ1/Δ1^* but not WT mice (Fig 5). In axonemal sections lacking normal RS assembly, irregular electron density was observed between the central and peripheral microtubules, potentially indicating an disorder of the RS complex. Partial RS absence was similarly detected in the spermatozoa of *Lrrc23^Δ2/Δ2^* mice (S7 Fig). A proteomic analysis revealed downregulation of RSPH3A/B and RSPH6A in the spermatozoa of *Lrrc23^Δ2/Δ2^* mice. Similarly, we observed downregulation of proteins associated with sperm motility including AKAP3, AKAP4 [27], and DNAH8 (Fig 6A), as confirmed via Western blotting (Fig 6B). However, no changes in levels of RSPH9 were detected in the spermatozoa of *Lrrc23^Δ1/Δ1^* males. High-resolution microscopy revealed a discontinuous or fragmented RSPH9 signal along the sperm flagella of *Lrrc23^Δ2/Δ2^* mice (Fig 6C), suggesting that the loss of flagellar LRRC23 resulted in RS structural defects.

**Fig 5.**
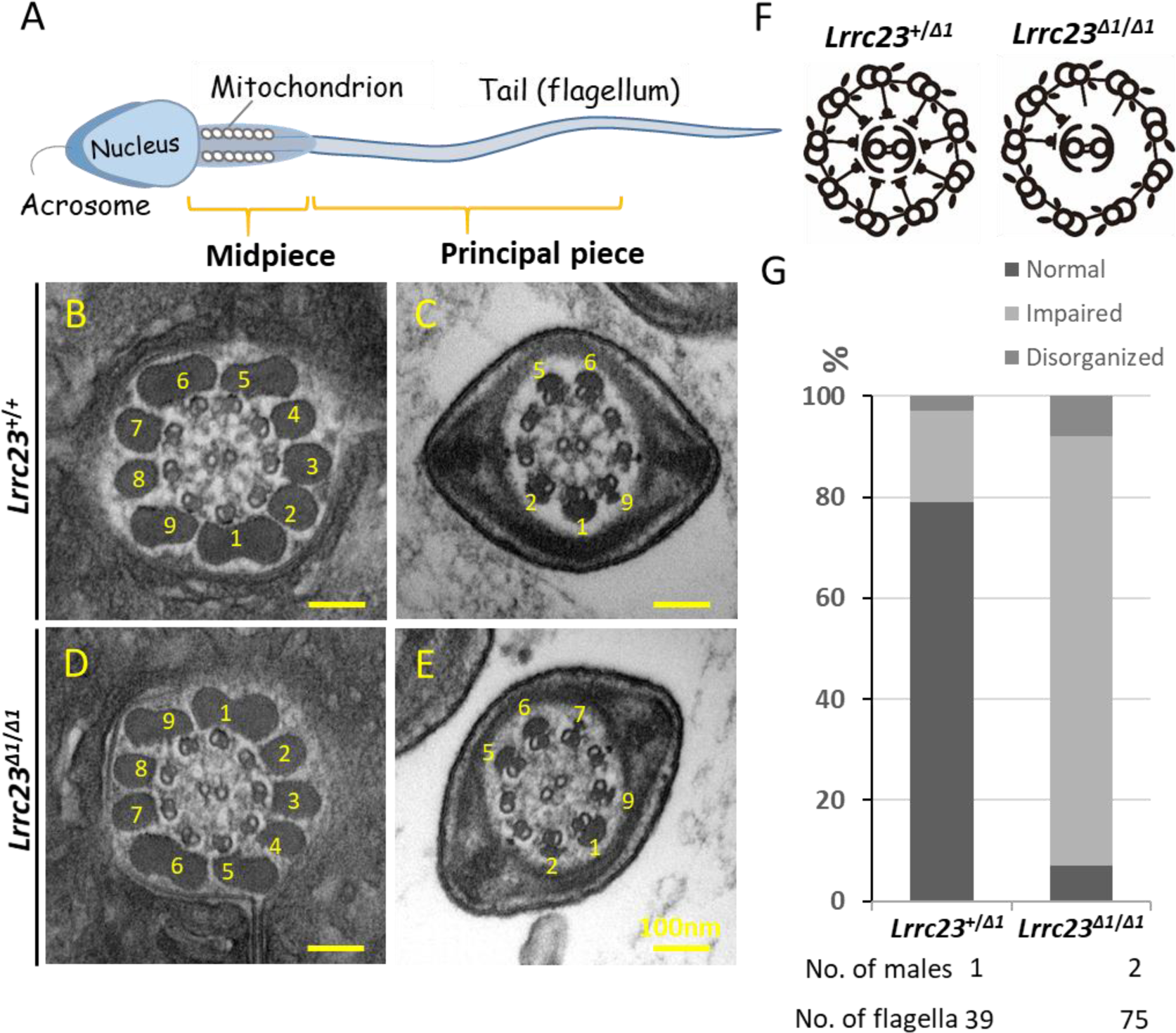
Ultrastructural analysis of cauda epididymis spermatozoa from *Lrrc23^Δ1/Δ1^* mice. (A) An overview of the structure of a mature spermatozoon. (B and C) Electron microscopy images demonstrating normal radial spoke structures in WT spermatozoa; (D and E) radial spokes were partially formed or absent in *Lrrc23^Δ1/Δ1^* sperm. Outer dense fibers are marked with numbers. (F) Schematic representation of the comparison between the axonemal structures of spermatozoa from *Lrrc23^+/+^* and *Lrrc23^Δ1/Δ1^* mice. (G) Percentages of the different axonemal structures in spermatozoa from *Lrrc23^+/+^* and *Lrrc23^Δ1/Δ1^* mice.

**Fig 6.**
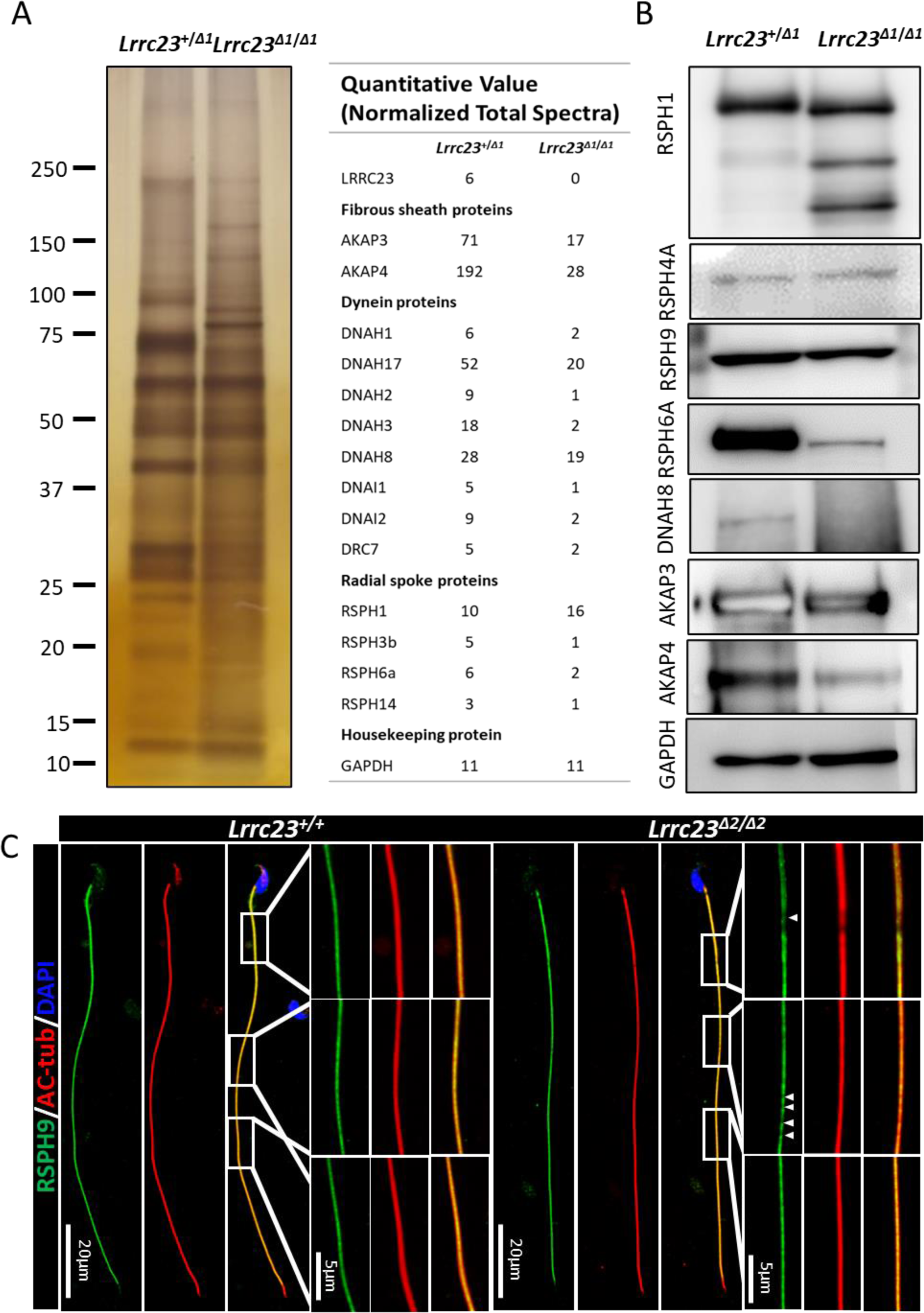
*Lrrc23* knockout causes the downregulation of other flagella component proteins. (A) Left: SDS-PAGE and silver staining were conducted to compare spermatozoa proteins in solubilized membrane fractions from *Lrrc23^+/+^* and *Lrrc23^Δ2/Δ2^* mice; Right: mass spectrometry revealed the downregulation of certain flagella components in the spermatozoa of *Lrrc23^Δ2/Δ2^* mice. (B) Western blotting was used to assess flagellum components in protein extracts from spermatozoa. (C) Immunofluorescence staining was conducted for spermatozoa obtained from WT and *Lrrc23^Δ2/Δ2^* males using antibodies specific for AC-tub (red) and RSPH9 (green). Nuclei were identified through Hoechst staining (blue). While rectangles indicate regions that have been magnified, with areas of RSPH9 signal discontinuity being marked with white arrows.

Since *Lrrc23* shows minor expression in lung, we investigated the functionality of the respiratory cillia in knockout mice by analyzing the expression of NME5, DYDC1, RSPH9, and HYDIN in tracheal cilia via high-resolution immunofluorescence microscopy, but no differences were observed in the expression or distributions of these proteins between the samples from *Lrrc23^Δ2/Δ2^* and *Lrrc23^+/+^* mice (Fig 7A). SEM analyses similarly uncovered obvious differences in respiratory cilia morphology in *Lrrc23^Δ2/Δ2^* and *Lrrc23^+/+^* animals (Fig 7B and 7C). Further, comparable tracheal ciliary beating was also detected in *Lrrc23^Δ2/Δ2^* and *Lrrc23^+/+^* mice (S3 Video, S4 Video), and no symptoms of epididymal ciliary abnormalities such as hydrocephaly, embryonic death, or poor postnatal survival were observed in *Lrrc23^Δ2/Δ2^* mice. Together, these data indicate that LRRC23 is an essential mediator of RS stability in mammalian sperm flagella, whereas it is dispensable for normal ciliary function.

**Fig 7.**
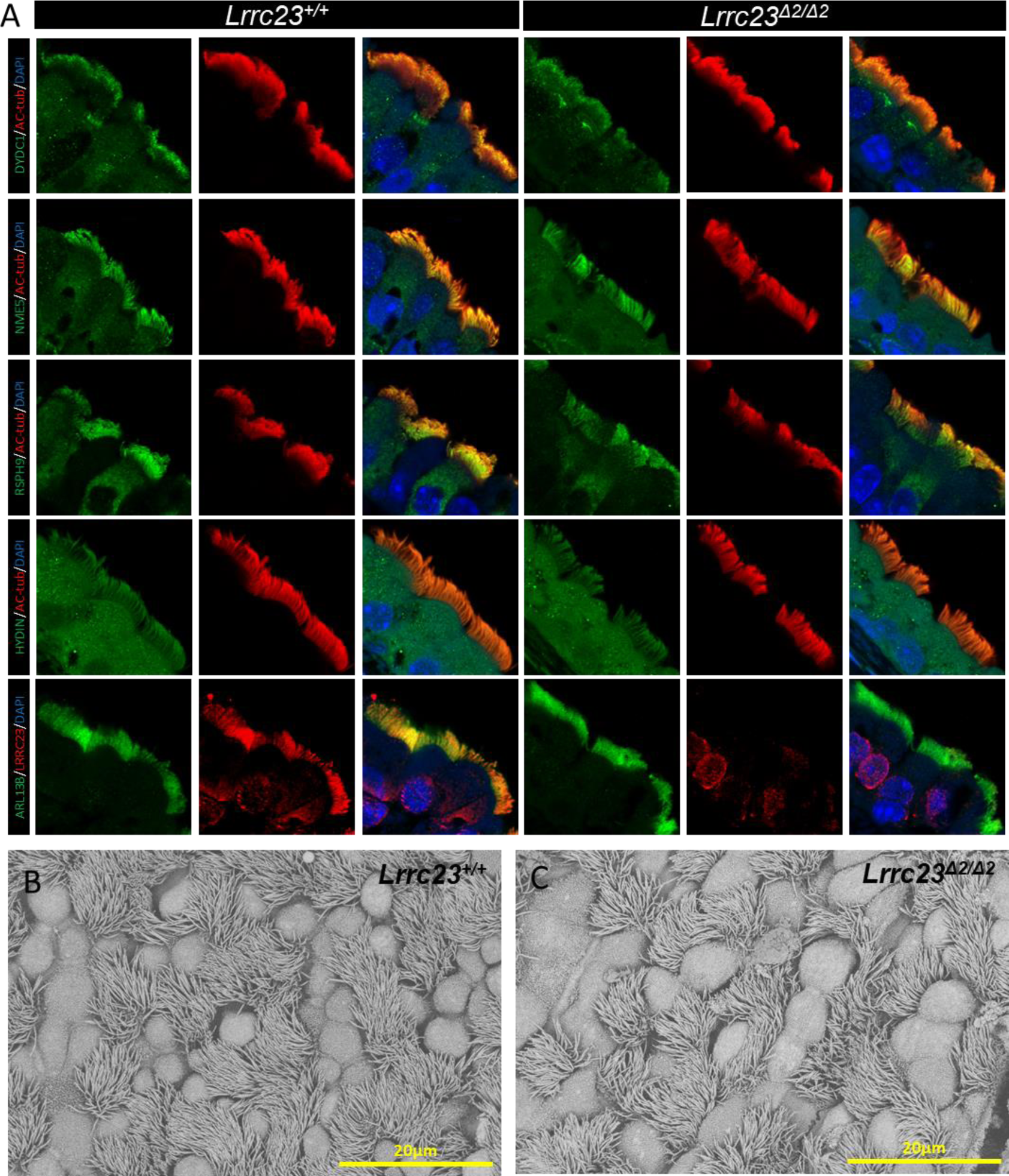
No significant differences in tracheal cilia components are evident when comparing *Lrrc23^+/+^* and *Lrrc23^Δ2/Δ2^* mice. (A) Immunofluorescent staining of respiratory tract cilia from the indicated mice was performed using antibodies specific for AC-tub, LRRC23 (red), DYDC1, NME5, RSPH9, ARL13B, and HYDIN (green). Nuclei were stained with Hoechst 33342 (blue). (B-C) Tracheal cilia from *Lrrc23^+/+^* and *Lrrc23^Δ2/Δ2^* mice were examined via SEM.

### Sterility of knockout males is rescued by transgenic expression of LRRC23

To determine whether the absence of *Lrrc23* was responsible for the male infertility, we produced a transgenic (Tg) mouse line expressing FLAG/1D4-tagged LRRC23 driven by the testicular germ cell-specific *Clgn* promoter on a *Lrrc23* knockout background [28] (Fig 8A). Western blot analysis confirmed that FLAG or 1D4-tagged LRRC23 was detected in both testis and sperm lysates, whereas no signal was detected in the testis or spermatozoa of WT males that did not carry the transgene (Fig 8B). *Lrrc23* knockout-Tg males were housed with WT females and resulted in normal litter sizes (8.9 ± 2.9, number of litters n = 152; Fig 8C), indicating that the knockout phenotype was rescued by the transgene. These results confirm that LRRC23 is required for normal sperm motility and male reproduction.

**Fig 8.**
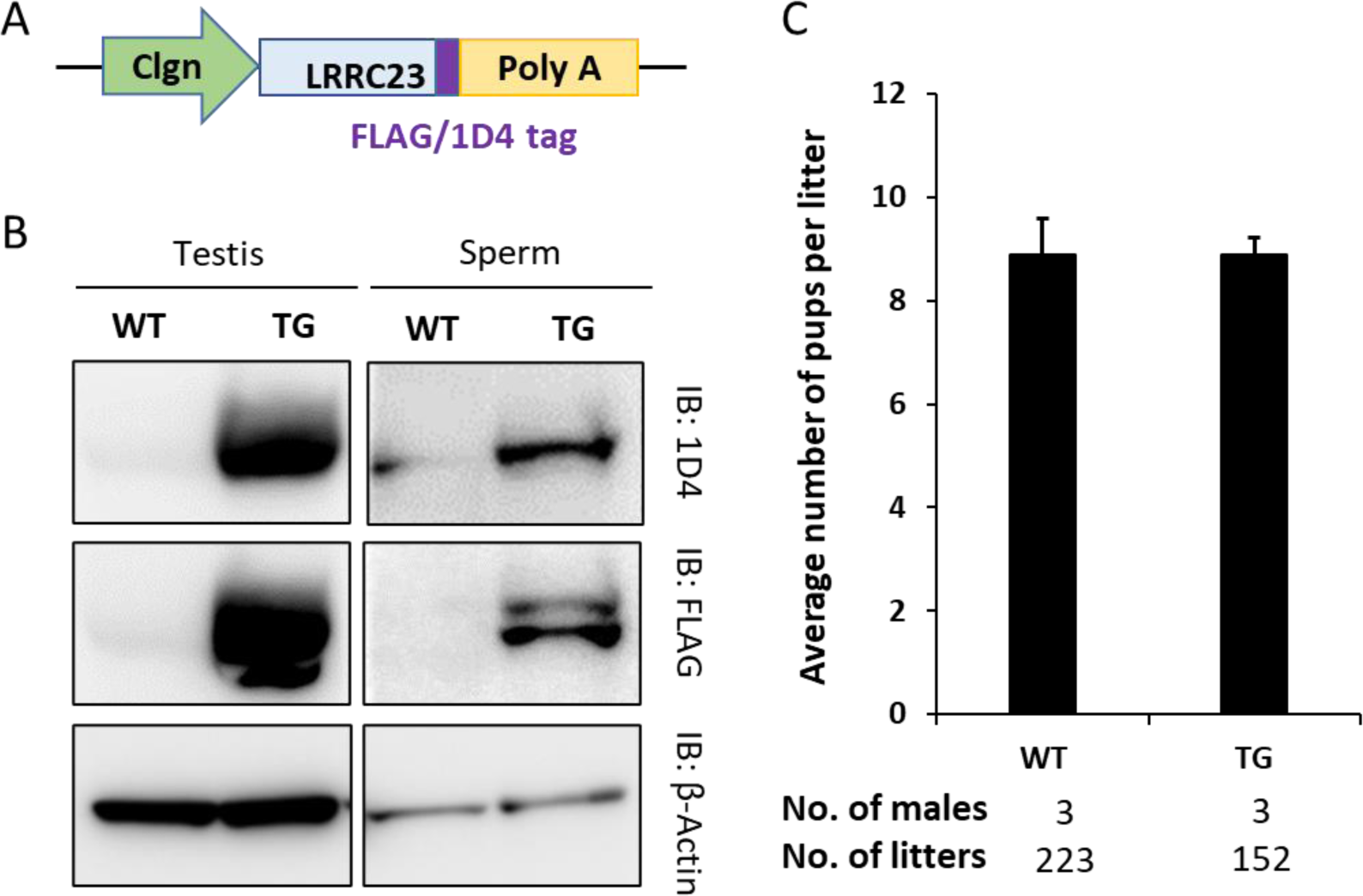
Sterility of KO males is rescued by transgenic expression of LRRC23. (A) Schematic representation of generating the *Lrrc23* transgene (Tg) mouse line. Transgene designed in which FLAG or 1D4-tagged LRRC23 are expressed as a fused protein under the testicular germ cell-specific *Clgn* promoter. (B) FLAG or 1D4-tagged LRRC23 expression in WT and knockout mice carrying the *Lrrc23* transgene was detected by Western blotting. β-Actin was analyzed as a loading control. (C) Average litter size of *Lrrc23* knockout male mice expressing FLAG or 1D4-tagged *Lrrc23* Tg. The average litter size (mean ± SD) was 8.9 ± 1.6 in WT males and 8.9 ± 2.9 in FLAG or 1D4-tagged *Lrrc23* rescued KO male mice.

## Discussion

Primary ciliary dyskinesia (PCD) is a genetic condition that arises as a consequence of ciliary and flagellar motility defects in multiple organ systems [29, 30], affecting the human race at a scale of 1:10,000 globally [31, 32]. While gross axonemal structures in sperm flagella and motile cilia appear similar, there are cell type-specific differences in axonemal assembly and utilized dynein arm components that differentiate these structures at the molecular level [33]. Herein, we report a distinct form of PCD. The depletion of LRRC23 led to impaired sperm motility but did not compromise respiratory ciliary motility. Light microscopy revealed that asthenospermia occurred in *Lrrc23* knockout mice, but no morphological abnormalities were observed. We have previously reported a similar form of PCD in *Tcte1* knockout mice [34]. While there have been no reports to date linked *LRRC23* or *TCTE1* mutations to asthenospermia in humans, we nonetheless believe that these findings would offer important insights into the tissue-specific differences in disease-related phenotypes that manifest in PCD patients.

Through high resolution fluorescence and immunoelectron microscopic analyses, we determined that LRRC23 localized near the center of the axonemal structure. Co-IP analyses further indicated that LRRC23 was able to interact with RSPH9, suggesting that the latter protein may also play a role in regulating the RS-related sperm flagellar functionality [35]. No significant abnormality in a 12S precursor RS component, RSPH9, was detected in cilia following *Lrrc23* knockout in mice, suggesting that this protein is not a component of the 12S precursor complex, consistent with findings observed for the homologous RSP13 protein in *Chlamydomonas* [36]. While RS abnormalities were detected in spermatozoa, no significant changes in the length of LRRC23-null sperm flagella were detected, in marked contrast to the short-tail phenotype observed in *Rsph6a* knockout spermatozoa. Given that the homologue of RSPH6A in *Chlamydomona*, RSP6, functions as a key 12S precursor complex component [36], these data indicate that the absence of 12S precursor components may pose a profound adverse impact on the flagellar formation. In contrast, the loss of LRRC23 exhibited compromised stability of sperm flagellar structures, such as dyneins, RSs and fibrous sheaths. Since dyneins serve as molecular motors and the RS complex functions as a mechanochemical sensor underpinning the flagellar motility [10], the disruption of these axonemal components can thus compromise the motility-related signal transmission and cause asthenospermia.

Through IP-MS analysis, we found that LRRC23 was able to interact with additional proteins beyond RSPH9, including SDF4, SSSCA1, PASK, and TDRD1. Since PASK, PAS domain-containing serine/threonine-protein kinase, was identified as a potential LRRC23-interacting protein, it is tempting to speculate that LRRC23 may be a kinase substrate activated by post-translational phosphorylation. While cAMP is known to be an important second messenger linked to sperm motility, our data did not support the existence of a cAMP-dependent protein kinase (PKA)-RSP3 regulatory pathway as has been reported in *Chlamydomonas* [37]. Whether the identified LRRC23-interacting proteins are necessary for sperm axonemal stability and/or flagellar beating remains to be determined.

Overall, we demonstrated LRRC23 as an RS protein that plays key roles in regulating sperm motility and sperm flagellar ultrastructural integrity. These data offer a theoretical basis for the incidence of asthenospermia and highlight novel targets that can be studied for better understanding of the mechanistic basis underlying sperm motility.

## Materials and methods

### Animals

All mice used in the present study were housed under standard conditions (20–22°C, 50-70% humidity, 12 h light/dark cycle) with free food and water access. The Institutional Animal Care and Use Committees of Nanjing Medical University and Osaka University approved all studies carried out by both laboratories (Approval No. IACUC-1810020; #Biken-AP-H30-01), and the Animal Ethical and Welfare Committee of both universities reviewed all animal protocols. Transgenic mice with Red Body Green Sperm [RBGS; B6D2-Tg(CAG/Su9-DsRed2, Acr3-EGFP)RBGS002Osb] prepared and housed in the laboratory of M.I were utilized to assess spermatozoa migration through the uterotubal junction (UTJ).

### In silico expression analysis

Murine testis transcriptome analyses were carried out using previous single-cell transcriptomic data [24]. *Lrrc23* mRNA levels in spermatogenic cells and related somatic cells were evaluated using the Loupe Cell Browser 3.3.1 (10X Genomics).

### RT-PCR

Total RNA was extracted from the tissues of adult ICR mice and from the testes of male mice from one to five weeks old, and a SuperScript III First Strand Synthesis Kit (Thermo Fisher, MA, USA) was used to prepare cDNA based on the manufacturer’s instructions. The expression of *Lrrc23* was assessed via PCR using the primers RT Fw and RT Rv as shown in Table S1.

### Generation of *Lrrc23^Δ1^* mutant mice by CRISPR/Cas9

The CRISPR/Cas9 system was used to generate *Lrrc23^Δ1^* mice. Two single-guide RNAs (sgRNAs; 5’-CATCATGGCCTCCGTGATGG -3’ and 5’-GGCTGGGCACACGGGACGAG -3’) were designed to target the exon 3 and exon 7 of *Lrrc23*, and their potential for off-target genomic editing was assessed using the CRISPRdirect program (crispr.dbcls.jp) [38]. Female B6D2F1 mice were superovulated via intraperitoneal injection with pregnant mare serum gonadotropin (PMSG) and human chorionic gonadotropin (hCG; ASKA Animal Health, Tokyo, Japan), after which they were paired with WT B6D2F1 males. The resultant two-pronuclear zygotes were isolated from the oviducts of the superovulated female mice, and a NEPA21 electroporation instrument (NEPA GENE, Chiba, Japan) was used to introduce crRNA/tracrRNA/Cas9 ribonucleoprotein complexes into the zygotes that were subsequently incubated in Potassium Simplex Optimized Medium (KSOM) medium [39] to the two-cell stage and transplanted into the ampullary segment of the oviducts in 0.5 d pseudopregnant ICR female mice. Founder animals were then obtained via natural delivery or Cesarean section after 19 d of pregnancy, with genotyping being conducted via Sanger sequencing and PCR. The genotyping primers are available in Table S1 (Fw#1, Rv#1 and Rv#2).

### Generation of *Lrrc23^Δ2^* mutant mice by CRISPR/Cas9

The CRISPR/Cas9 system was used to generate *Lrrc23^Δ2^* knockout mice using two sgRNAs targeted to knockout the exon 4 of *Lrrc23* (5’-GCAATCAGCTTCGGAGTGCT -3’ and 5’-GATCTGGTTGTAGGAAAAGC -3’). Two complementary DNA oligos for each of these sgRNA targets were annealed and ligated to the BsaI-digested pUC57-T7-sgRNA vector, while sgRNA templates were amplified from sgRNA plasmids via PCR using primers (Fw trans and Rv trans, shown in Table S1). A MinElute PCR Purification Kit (QIAGEN, Duesseldorf, Germany) was then used to isolate the amplified template sequences, and sgRNAs were generated with the MEGAshortscript Kit (Ambion, Austin, TX, USA) and purified with a MEGAclear Kit (Ambion, Austin, TX, USA) based on provided directions. Following linearization with AgeI, a Cas9 plasmid (Addgene, Watertown, MA, USA) was purified using a MinElute PCR Purification Kit (QIAGEN, Duesseldorf, Germany), after which an mMESSAGE mMACHINE T7 Ultra Kit (Ambion, Austin, TX, USA) was used to transcribe Cas9 mRNA that was subsequently purified with an RNeasy Mini Kit (QIAGEN, Duesseldorf, Germany) based upon provided directions. The Cas9 mRNA (50 ng/μL) and sgRNA (20 ng/μL) were then co-injected into murine zygotes which were transferred into pseudopregnant females. On postnatal day 7, toe clipping was conducted to tag the newborn mice, and DNA was extracted from these tissue samples with a Mouse Direct PCR Kit (Biotool, Shanghai, China). Sanger sequencing was performed after PCR amplification with appropriate primers (Fw#3 and Rv#4, S1 Table) and PrimeSTAR HS DNA Polymerase (Takara, Kyoto, Japan).

### Fertility Testing

Three *Lrrc23^Δ1/Δ1^* and three *Lrrc23^Δ2/Δ2^* male sexually mature mice were individually housed with three 8-week-old WT B6D2F1 female mice for a minimum of 2 months, with WT male mice undergoing the same housing conditions as a control. Litter sizes were recorded at the date of birth, with average litter size being calculated by dividing total numbers of pups by total numbers of litters.

### Antibody preparation

Antibodies were produced as detailed previously [40]. The full-length murine *Lrrc23* cDNA (aa 1–340) was expressed as His fusion protein in *E. coli* with the pET-28a(+) vector, and the Ni-NTA His Bind Resin (TransGen Biotech, Beijing, China) was used to achieve the affinity purification of the resultant protein which was then used to immunize two rabbits were immunized with the fusion protein, yielding anti-LRRC23 antisera.

Monoclonal antibody of LRRC23 used for Immunogold-EM and sperm protein fractionation analyses was generated as previously described. The sequence encoding mouse LRRC23 (residue 171-340 aa, CCDS20528.1) was cloned and inserted into pGEX6p-1 expression vector (GE healthcare), followed by transformation into *E. coli* strain BL21 (de3) pLysS (C606003, Thermo Fisher Scientific, USA). GST-fused LRRC23 recombinant protein was expressed and subjected to the treatment of PreScission Protease to remove the GST tag [41]. The recombinant LRRC23 protein was then purified and injected into female rats in combination with a complete adjuvant. After 17 days of injection, lymphocytes were collected from iliac lymph nodes and hybridomas were generated and cultured [42]. The supernatants obtained from the hybridomas were used as antibodies. The candidates were screened by ELISA against LRRC23.

### Testis weights and sperm motility analyses

*Lrrc23* knockout males were anesthetized and euthanized via cervical location, after which testis and body weight were measured for both heterozygous and homozygous knockout animals. Spermatozoa were extracted from cauda epididymides, after which the motility of these cells was assessed with a CEROS II Computer-assisted sperm analysis (CASA) system (Hamilton Thorne Biosciences, MA) at 10 min and 2 h following incubation in Toyoda, Yokoyama, Hoshi (TYH) medium. Sperm movement was additionally recorded at 200 frames/second using an Olympus BX-53 microscope equipped with a high-speed camera (HAS-L1, Ditect, Tokyo, Japan). The BohBoh sperm motion analysis software was then used to reconstruct sperm flagellar waveforms based on these videos (BohBohsoft, Tokyo, Japan).

After extraction, sperm derived from the epididymis of *Lrrc23****^Δ2/Δ2^*** mice or littermate controls (*Lrrc23****^Δ2/+^*** or WT males) were incubated in human tubal fluid (HTF) media (FUJIFILM Irvine Scientific, Japan) containing 10% FBS at 37°C, with Hamilton Thorne’s Ceros II system (Hamilton-Thorne Research, Inc., Beverly, MA, USA) being used to dilute and analyze these samples. For PKA inhibition experiments, sperm were incubated in HTF medium containing 10% FBS in the presence or absence of H-89 or BioE-1115 (S3 Table) for 15 min in air at 37 °C. After 15 min at 37 °C, these samples were analyzed via CASA system and western blotting.

### Sperm UTJ migration assay

WT B6D2F1 females were superovulated by peritoneal injection of pregnant mare serum gonadotropin (PMSG) and hCG. After 12 h of hCG injection, the hormone primed females were individually housed with a Lrrc23 knockout male mouse expressing DsRed2/Acr3-EGFP. Success of copulation was confirmed by the formation of a vaginal plug. The female mice were sacrificed 2 h after copulation and the female reproductive tract was collected for imaging. Spermatozoa with red fluorescence in the midpiece and green fluorescence in the acrosome were observed directly under a Keyence BZ-X710 microscope (Keyence, Osaka, Japan).

### Analyses of testis and epididymis histology and sperm morphology

Testes and epididymides were fixed in Bouin’s solution (Polysciences Inc., Warrington, PA) and embedded in paraffin wax. Paraffin sections were stained with periodic acid (Nacalai Tesque, Kyoto, Japan) and Schiff’s reagent (Wako, Osaka, Japan) and counterstained with Mayer hematoxylin solution (Wako, Osaka, Japan). The cauda epididymal spermatozoa were dispersed in the TYH medium and observed under an Olympus BX53 phase contrast microscopy (Olympus, Tokyo, Japan).

### Transmission electron microscopy (TEM)

Ultrastructural analyses of testes and spermatozoa using TEM were conducted as previously described [43]. Male WT and *Lrrc23^Δ1/Δ1^* mice were anesthetized and perfused with 4% paraformaldehyde (PFA), after which fixed cauda epididymis tissues were collected and fixed for an additional 6 h in 4% PFA at 4°C. These epididymides were then minced with razor blades to yield 2 mm cubes that were fixed overnight with 1% glutaraldehyde in 30 mM HEPES (pH 7.8) at 4°C. Post-fixation for 1 h with 1% osmium tetroxide (OsO_4_) and 0.5% potassium ferrocyanide in 30 mM HEPES was then performed at room temperature, after which an ethanol gradient was used to dehydrate samples, which were then embedded for 2 days using epoxy resin at 60°C for 2 days. An Ultracut Microtome was then used to prepare ultrathin sections that were stained with both uranyl acetate and lead citrate prior to mounting onto copper grids and evaluation with a JEM-1400 Plus electron microscope (JEOL, Tokyo, Japan) at 80 kV equipped with a Veleta 2k × 2k CCD camera (Olympus, Tokyo, Japan).

Similarly, spermatozoa from *Lrrc23^Δ2/Δ2^* male mice were fixed overnight with 2.5% glutaraldehyde, post-fixed with 2% OsO_4_, and embedded in Araldite for ultrastructural analyses. Ultrathin (80 nm) sections were then stained using uranyl acetate and lead citrate and were imaged via EM (JEM.1010, JEOL).

### Scanning electron microscopy (SEM)

Spermatozoa samples were fixed for 2 h with 2.5% phosphate-buffered glutaraldehyde at 4°C. Spermatazoa were then allowed to attach to coverslips coated with poly-L-lysine. Both sample types were then washed with PBS, dehydrated with a chilled ethanol gradient (30%, 50%, 70%, 80%, 90%, and 100%), and subjected to critical point drying with a Lecia EM CPD300 Critical Point Dryer (Wetzlar, Germany). Samples were then attached to appropriate specimen holders and coated with gold particles via the use of an ion sputter coater (EM ACE200, Leica). A Helios G4 CX scanning electron microscope (Thermo Scientific) was then used to image samples.

### In vitro fertilization (IVF)

Spermatozoa isolated from the cauda epididymides of WT and *Lrrc23* knockout male mice were suspended in TYH medium [44]. Cumulus-oocyte complexes (COCs) were collected from superovulated B6D2F1 female mice, and were treated for 5 min with 1.0 mg/ml hyaluronidase (Wako, Osaka, Japan) at 37°C to remove the cumulus cells, or with 1.0 mg/ml collagenase (Sigma, St. Louis, MO) to remove the zona pellucida (ZP). Cumulus-intact and cumulus-free oocytes were then inseminated by combing them with 2.0 × 10^5^ sperm/mL, while ZP-free oocytes were combined with 2.0 × 10^4^ sperm/mL. Following a 6 h insemination period, the formation of two pronuclei was assessed to gauge fertilization success.

### Sperm protein fractionation

The fractionation of sperm proteins was performed as in prior reports [45]. Briefly, 1% Triton X-100 lysis buffer (50 mM NaCl, 20 mM Tris-HCl, pH 7.5) was used to lyse isolated spermatozoa for 2 h at 4°C. Supernatants containing the Triton-soluble fraction were then collected after spinning for 10 min at 15,000 × *g*, while the insoluble pellets were subjected to lysis for 1 h with 1% SDS lysis buffer (75 mM NaCl, 24 mM EDTA, pH 6.0) at room temperature. After being spun down for an additional 10 min at 15,000 × *g*, the SDS-soluble supernatant fraction was collected, while sample buffer (66 mM Tris-HCl, 2% SDS, 10% glycerol, and 0.005% Bromophenol Blue) was used to lyse the SDS-resistant pellet, and samples were boiled for 5 min, with the SDS-resistant fraction being isolated following centrifugation.

### Western blotting

Urea/thiourea lysis buffer (8 M urea, 50mM Tris-HCl pH 8.2, 75Mm NaCl) containing a 2% (v/w) protease inhibitor cocktail (Roche, Basel, Switzerland) was used to extract proteins from murine tissue samples. These proteins were then separated via SDS-PAGE and transferred to PVDF membranes that were blocked for 2 h with 5% non-fat milk in TBST at room temperature, following by overnight incubation with primary antibodies. These primary antibodies are shown in S2 Table. The membranes were then washed thrice in TBST, probed for 2 h with appropriate secondary antibodies (S2 Table), and High-sig ECL Western Blotting Substrate (Tanon, Shanghai, China) Western Blotting Detection system was then used to detect protein bands. For phosphatase treatment experiments, the fractionation of sperm or testis proteins were at 1 mg/ml in CIP buffer (5mM Tris pH 8.2, 10mM NaCl, 1mM MgCl_2_, 0.1 mM DTT) and then incubating with 1.5 U of calf intestinal alkaline phosphatase (MilliporeSigma, Stockholm, Sweden) per 100 μg of protein at 37°C for 3h. The reaction was terminated by the addition of protein loading buffer and treated at 95°C for 10 min. The samples were resolved by 10% SDS-PAGE and analyzed by western blotting. Gray value was analyzed via ImageJ software (ImageJ 1.52a, USA)

### Immunoprecipitation

RIPA buffer (Beyotime, Shanghai, China) containing a 2% proteinase inhibitor cocktail was used to lyse samples at 4°C for 40 min, after which samples were spun down for 40 min at 12,000 × *g*. Supernatants were then mixed for 1 h with Protein A magnetic beads (Thermo Fisher Scientific, MA, USA), and lysates were then incubated overnight with primary anti-LRRC23 or Rabbit IgG (#2729, Cell Signaling Technology, Danvers, Massachusetts, USA) at 4°C. Samples were then mixed for 3 h with 50 μL of Protein A magnetic beads at 4°C, after which they were washed with PBST. IP pellets and extract samples were then divided in two, with half being used for western blotting after boiling in SDS loading buffer and half being subjected to mass spectrum analyses performed by Genedenovo Biotechnology (Guangzhou, China).

### Immunofluorescent staining

The immunofluorescent staining of tissue sections was conducted as detailed previously [46]. For analyses of sperm cells, these samples were smeared onto slides, air-dried, fixed for 40 min with 4% paraformaldehyde, washed thrice with PBS (5 min/wash), and antigen retrieval was then conducted by boiling slides in a microwave for 10 min in 10 mM citrate buffer (pH 6.0). Following three additional washes in PBST (PBS containing 0.05% Tween-20; 10 min/wash), 5% BSA in PBST was used to block slides for 2 h, after which they were probed overnight with appropriate primary antibodies (Table S2) at 4°C. After three additional PBST washes, slides were probed for 2 h with secondary antibodies (S2 Table), counterstained for 5 min with Hoechst 33342 (Invitrogen, Carlsbad, CA, USA), rinsed with PBST, mounted, and imaged with a LSM800 confocal microscope (Carl Zeiss AG, Jena, Germany) and TCS SP8X confocal microscope (Leica Microsystems, Wetzlar, Germany).

### Immunoelectron microscopy

Cauda epididymides from WT and *Lrrc23****^Δ1/Δ1^*** mice were dissected after perfusion fixation of the whole bodies with 4% PFA in 0.1 M phosphate buffer under anesthesia. The epididymides were then sliced into 3-4 mm thick sections and fixed in 4% formaldehyde in 0.1 M phosphate buffer (pH 7.4) for 1 h at room temperature. After fixation, the samples were washed with 4% sucrose in 0.1 M phosphate buffer (pH 7.4) for three times. Tissue samples were then incubated in 10%, 15%, and 20% sucrose in 0.1 M phosphate buffer (pH 7.4) sequentially for 6 h each and embedded in OCT compound (Sakura, Tokyo, Japan), and frozen in liquid nitrogen. The samples were subsequently sectioned under −20 °C using a cryostat (Thermo), and the cryo-sections were attached to MAS coated glass coverslips (Matsunami Glass) and air-dried for 30 min. The coverslips were placed in 24-well culture plate and blocked with blocking solution (0.1M phosphate buffer containing 0.1% saponin, 10% BSA, 10% normal goat serum and 0.1% cold water fish skin gelatin) for 30 min. Gold particle labeling was performed using primary antibody, Rat anti-LRRC23 antibody 1:150 in blocking solution and second antibody, goat anti-rat IgG coupled to 1.4 nm gold 1:300 (Nanogold, Nanoprobes, Yaphank, NY, USA) in blocking solution followed the procedures as previously described [47]. After post-fixed in 1% OsO_4_ and 1.5% potassium ferrocyanide in 0.1 M phosphate buffer (pH 7.4) for 1 h, samples were dehydrated in a graded series of ethanol, substituted with propylene oxide, and embedded in epoxy resin. Ultrathin sections were stained with 8% uranyl acetate and lead staining solution. The samples were examined using a JEM-1400 plus electron microscope (JEOL, Tokyo, Japan) at 80 kV with a CCD Veleta 2K × 2K camera (Olympus).

### Mass spectrometry

LRRC23 was immunoprecipitated from mouse testis using the Pierce crosslink IP kit (Thermo Scientific, MA, USA) with anti-LRRC23 antibody described above, IgG antibody was used as negative control. IP was performed according to the manufacturer’s instructions. Eluates were precipitated with five volumes of −20°C pre-chilled acetone followed by trypsin digestion. LC-MS/MS analysis was performed on EASY-nanoLC 1000 system (Thermo Scientific, MA, USA) coupled to an Orbitrap Fusion Tribrid mass spectrometer (Thermo Scientific, MA, USA) by a nano spray ion source. Tryptic peptide mixtures were injected automatically and loaded at a flow rate of 20 μl/min in 0.1% formic acid in LC-grade water onto an analytical column (Acclaim PepMap C18, 75 μm x 25 cm; Thermo Scientific). The peptide mixture was separated by a linear gradient from 5% to 38% of buffer B ( 0.1% formic acid in ACN) at a flow rate of 300 nl/min over 53 minutes. Remaining peptides were eluted by a short gradient from 38% to 90% buffer B in 1 minutes. Analysis of the eluted peptides was done on an Orbitrap Fusion Tribrid mass spectrometer. From the high-resolution MS pre-scan with a mass range of 335 to 1400, the most intense peptide ions were selected for fragment analysis in the orbitrap depending by using a high speed method if they were at least doubly charged. The normalized collision energy for HCD was set to a value of 28 and the resulting fragments were detected with a resolution of 120,000. The lock mass option was activated; the background signal with a mass of 445.12003 was used as lock mass. Every ion selected for fragmentation was excluded for 30 seconds by dynamic exclusion. Data were processed with MaxQuant software (version 1.6.10.43) and Mouse reference proteome from SwissProt database (release 2019_07) using standard parameters. LFQ was used as the main parameter for protein quantification [48]. In the control group, LFQ intensity should be 0 and no more than one group of samples should have peptide detection. The mass spectrometry proteomics data have been deposited to the ProteomeXchange Consortium via the PRIDE partner repository [49] with the dataset identifier PXD025549.

Whole sperm proteomic analyses were performed as previously described [50]. Briefly, protein samples were extracted from spermatozoa using lysis buffer (6 M urea, 2 M thiourea, and 2% sodium deoxycholate) and centrifuged at 15,000 × *g* for 15 min at 4°C. The samples were processed and the resultant protein peptides were subjected to nanocapillary reversed-phase LC-MS/MS analysis using a C18 column (10 cm x 75 um, 1.9 µm, Bruker Daltoniks, Bremen, Germany) on a nanoLC system (Bruker Daltoniks, Bremen, Germany) connected to a timsTOF Pro mass spectrometer (Bruker Daltoniks) and a nano-electrospray ion source (CaptiveSpray, Bruker Daltoniks). The resulting data was processed using DataAnalysis (Bruker Daltoniks), and proteins were identified using MASCOT Sever (Matrix Science, London, UK) against the SwissProt database. Quantitative value and fold exchange were calculated by Scaffold4 (Proteome Software, Portland, OR, USA). The raw data is accessible from the ProteomeXchange Consortium via the dataset identifier PXD025166.

### Production of transgenic mice

Fertilized eggs were obtained from in vitro fertilization between spermatozoa from *Lrrc23* heterozygous males and oocytes from homozygous or heterozygous knockout females. Linearized plasmids encoding a *Calmergin* (*Clgn*) promoter, a rabbit beta-globin polyadenylation signal, and FLAG or 1D4-tagged LRRC23 was microinjected into the pronuclei of the zygotes. The treated zygotes were then cultured in KSOM medium to two-cell stage and transplanted into the ampullary segment of the oviducts in 0.5 d pseudopregnant ICR females. Founder animals were obtained by natural delivery or Cesarean section after 19 d of pregnancy. Primer sequences used for genotyping of transgenic mice are enumerated in Table S1 (Fw#4, FLAG and 1D4).

## Acknowledgments

This work was supported by the National Key Research and Development Program of China 2016YFA0500902 (to M.L.); Natural Science Foundation of China (31771654 and 32070842 to M.L.); the Natural Science Foundation of Jiangsu Province (Grants No. BK20190081 to M.L.); and Qing Lan Project (to M.L.); Ministry of Education, Culture, Sports, Science and Technology/Japan Society for the Promotion of Science KAKENHI (Grants-in-Aid for Scientific Research) Grants JP18K16735 (to Y.L.); JP20K16107 (to K.S.); JP18K14612 and JP20H03172 (to T.N.); JP17H01394 and JP19H05750 (to M.I.); Takeda Science Foundation grants (to T.N. and M.I.). Japan Agency for Medical Research and Development Grant JP20gm5010001 (to M.I.); Eunice Kennedy Shriver National Institute of Child Health and Human Development Grants R01HD088412 and P01HD087157 (to M.I.); and the Bill & Melinda Gates Foundation Grant INV-001902 (to M.I.).

## Author contributions

M. I. and M.-X.L. initiated the project and designed the experiments; X.Z., J.S., Y.-G. L. and J.-T.Z. performed most of the experiments and analysis; K.S., T.N., S.-Q.Z., T.K., M.M., S.-S.Z., and J.-Y.W. performed some of the experiments and analysis; M.-X.L., X.Z., Y.-G. L. and J.S. wrote the manuscript; all authors read and approved the final manuscript.

## Statistical analysis

Data are given as mean ± SEM and were compared via two-tailed paired Student’s t-tests. * *P* < 0.05, ** *P* < 0.01, *** *P* < 0.001.

## Supporting information

**S1 Fig**. ***Lrrc23* is an evolutionarily conserved gene.** (A) *Lrrc23* is present in the majority of eukaryotes that utilize flagella (SAR: stramenopiles, alveolates, Rhizaria). Green denotes *Lrrc23* by the majority of species within the indicated taxon, whereas yellow indicates a loss of this gene within several species within the indicated taxon. (B) LRRC23 protein sequence similarity across species, with dark blue corresponding to total conservation and light blue indicating conservation among three to five species.

**S2 Fig**. ***In silico* analysis of *Lrrc23* expression in human and murine testes.** (A-B) *Lrrc23* is expressed in spermatocytes and round spermatids in murine testes, (C-D) while in human testis expression of this gene is evident in spermatogonia, spermatocytes, and round spermatids. Ud Spg: undifferentiated spermatogonia, D Sg: differentiated spermatogonia, A1-A2 Sg: A1-A2 differentiating spermatogonia, A3-B Sg: A3-A4-In-B differentiating spermatogonia, PreLe Sc: Preleptotene spermatocytes, Le/Zy Sc: Leptotene/Zygotene spermatocytes, Pa Sc: Pachytene spermatocytes, Di/Se Sc: Diplotene/Secondary spermatocytes, Early St: Early round spermatids, Mid St: Mid round spermatids, Late St: Late round spermatids, SC: Sertoli cells, PTM: Peritubular myoid cells, LC: Leydig cells, EC: Endothelial cells, PC: Perivascular cells

**S3 Fig. Assessment of the fertility of spermatozoa from *Lrrc23^Δ1/Δ1^*mice.** (A) Fertilization rates (percentages of two pronuclei [2PN] eggs) in cumulus-intact oocytes inseminated with spermatozoa from *Lrrc23^+/Δ1^* and *Lrrc23^Δ1/Δ1^* mice, N = 3, *P* < 0.001. (B) Fertilization rates in cumulus-free oocytes generated with spermatozoa from *Lrrc23^+/Δ1^* and *Lrrc23^Δ1/Δ1^* mice, N = 3, *P* < 0.05. (C) Fertilization rates in ZP-free oocytes generated with spermatozoa from *Lrrc23^+/Δ1^* and *Lrrc23^Δ1/Δ1^* mice, N = 3, *P* > 0.05.

**S4 Fig. Generation and analysis of male *Lrrc23^Δ2/Δ2^* mice.** (A) Dual sgRNAs (sgRNA#3 and sgRNA#4) were used to target *Lrrc23* exon 4, with Sanger sequencing being used to confirm the successful deletion of a 49 bp fragment within this region. Black rectangles are used to denote the coding regions, and genotyping primers (Fw#3, Rv#3) were as shown. (B) Testes of *Lrrc23^+/+^* and *Lrrc23^Δ2/Δ2^* mice. (C) Average testis weight/body weight in *Lrrc23^+/+^* and *Lrrc23^Δ2/Δ2^* mice, N = 3, *P* > 0.05. (D) Cauda epididymal sperm contents from *Lrrc23^+/+^* and *Lrrc23^Δ2/Δ2^* mice, N = 3, *P* > 0.05. (E) Normal epididymal sperm counts from *Lrrc23^+/+^* and *Lrrc23^Δ2/Δ2^* mice, N = 3, *P* > 0.05; (F) Average numbers of pups per litter from *Lrrc23^+/+^* and *Lrrc23^Δ2/Δ2^* mice, N = 3, *P* < 0.001. (G) Spermatozoa from *Lrrc23^+/+^* and *Lrrc23^Δ2/Δ2^* mice were subjected to hematoxylin and eosin staining. (H) SEM was used to image WT and *Lrrc23* knockout spermatozoa. (I) average percentages of motile spermatozoa and (J) progressively motile spermatozoa from *Lrrc23^+/+^* and *Lrrc23^Δ2/Δ2^* mice were quantified, N = 3, *P* < 0.05. (K) Flagellar waveforms for spermatozoa from *Lrrc23^+/+^* and *Lrrc23^Δ2/Δ2^* mice were assessed following a 5 min incubation.

**S5 Fig. LRRC23 is a radial spoke complex component that interacts with other proteins within this complex.** (A) LRRC23 immunoprecipitation (IP) in testicular protein extracts from *Lrrc23^+/+^* mice. (B, C, and D) Co-immunoprecipitation of LRRC23-FLAG and RSPH-HA was conducted using anti-FLAG-conjugated beads to examine interactions between these two proteins. Input: whole cell lysates from experimental cells; IP: samples immunoprecipitated with anti-FLAG beads. In HEK293T cells, LRRC23 was able to interact with other RS proteins including RSPH22 (B), RSPH3A (C), and RSPH3B (D).

**S6 Fig. LRRC23 interacts with PASK and undergoes phosphorylation in testis and spermatozoa.** (A) Co-immunoprecipitation of LRRC23-FLAG and PASK-HA was conducted using anti-FLAG-conjugated beads to examine interactions between these two proteins. Input: whole cell lysates from experimental cells; IP: samples immunoprecipitated with anti-FLAG beads. (B) Testis and sperm proteins from wild-type mice were treated with CIP. Or were left untreated (Con), revealing the phosphorylation of LRRC23 in control samples. Densitometric values along the vertical axis in the indicated region are shown to the right, with numbers corresponding to individual lanes and with β-tubulin serving as a control. (C, D, and E) WT spermatozoa were treated with inhibitors specific for cyclic AMP-dependent protein kinase (H-89) and PASK (BioE-1115) for 5 min, after which no significant LRRC23 phosphorylation was detectable (C). (D) H-89 treatment reduced sperm motility, whereas (E) BioE-1115 had no impact on this motility.

**S7 Fig. Ultrastructural assessment of spermatozoa in the cauda epididymis of *Lrrc23^Δ2/Δ2^* mice.** (A and B) Electron microscopy was used to assess cross sections of the principal component of spermatozoa from *Lrrc23^+/+^* and *Lrrc23^Δ2/Δ2^* mice. Outer dense fibers are marked with numbers, while the absence of a radial spoke is marked by red arrows.

**S1 Video**. **Spermatozoa from *Lrrc23^+/Δ1^* mice.** Spermatozoa of *Lrrc23^+/Δ1^* mice at 10 min of incubation in TYH media. Movie is recorded at 200 frames/second using an Olympus BX-53 microscope equipped with a high-speed camera (HAS-L1, Ditect, Tokyo, Japan).

**S2 Video**. **Spermatozoa from *Lrrc23^Δ1/Δ1^* mice.** Spermatozoa of *Lrrc23* knockout mice at 10 min of incubation in TYH media. Movie is recorded at 200 frames/second using an Olympus BX-53 microscope equipped with a high-speed camera (HAS-L1, Ditect, Tokyo, Japan).

**S3 Video. The beating of respiratory cilia in *Lrrc23^+/Δ2^* mice.**

**S4 Video. The beating of respiratory cilia in *Lrrc23^Δ2/Δ2^* mice.**

**S1 Table. Primer sequences.**

**S2 Table. List of Antibodies.**

**S3 Table. List of Inhibitors.**

